# Cell size-dependent regulation of Wee1 localization by Cdr2 cortical nodes

**DOI:** 10.1101/195628

**Authors:** Corey A. H. Allard, Hannah E. Opalko, Ko-Wei Liu, Uche Medoh, James B. Moseley

## Abstract

Cell size control requires mechanisms that link cell growth with Cdk1 activity. In fission yeast, the protein kinase Cdr2 forms cortical nodes that include the Cdk1 inhibitor Wee1, along with the Wee1-inhibitory kinase Cdr1. We investigated how nodes inhibit Wee1 during cell growth. Biochemical fractionation revealed that Cdr2 nodes were megadalton structures enriched for activated Cdr2, which increases in level during interphase growth. In live-cell TIRF movies, Cdr2 and Cdr1 remained constant at nodes over time, but Wee1 localized to nodes in short bursts. Recruitment of Wee1 to nodes required Cdr2 kinase activity and the noncatalytic N-terminus of Wee1. Bursts of Wee1 localization to nodes increased 20-fold as cells doubled in size throughout G2. Size-dependent signaling was due in part to the Cdr2 inhibitor Pom1, which suppressed Wee1 node bursts in small cells. Thus, increasing Cdr2 activity during cell growth promotes Wee1 localization to nodes, where inhibitory phosphorylation of Wee1 by Cdr1 and Cdr2 kinases promotes mitotic entry.

**Summary:** Cells turn off the mitotic inhibitor Wee1 to enter into mitosis. This study shows how cell growth progressively inhibits fission yeast Wee1 through dynamic bursts of localization to cortical node structures that contain Wee1 inhibitory kinases.

## Introduction

Many cell types divide at a reproducible size due to poorly understood mechanisms that couple cell growth to cell cycle signaling (Dolznig et al., 2004; Ginzberg et al., 2015). In eukaryotes, the ubiquitous cyclin-dependent kinase Cdk1 triggers mitotic entry and cell division (Harashima et al., 2013). During G2, the protein kinase Wee1 phosphorylates and inhibits Cdk1 to prevent premature mitosis (Gould and Nurse, 1989; Russell and Nurse, 1987). The counteracting phosphatase Cdc25 removes this inhibitory phosphorylation to activate Cdk1 and promote mitotic entry (Gautier et al., 1991; Kumagai and Dunphy, 1991; Russell and Nurse, 1986; Strausfeld et al., 1991). The balance of Wee1 versus Cdc25 activity determines the timing of mitotic entry and cell division, meaning that cells require mechanisms to inhibit Wee1 and activate Cdc25 as they grow during G2 (Moreno et al., 1989).

This conserved Cdk1 activation system was initially identified and characterized in the fission yeast *Schizosaccharomyces pombe* (Gould and Nurse, 1989; Russell and Nurse, 1986, 1987; Simanis and Nurse, 1986). These rod-shaped cells grow by linear extension at the cell tips with no change in cell width, and then enter mitosis and divide at a threshold size due to the regulated activation of Cdk1 (Fantes and Nurse, 1977; Moreno et al., 1989). The concentration of Cdc25 protein increases as cells grow in G2, providing a simple mechanism for its size-dependent regulation (Keifenheim et al., 2017; Moreno et al., 1990). In contrast, the concentration of Wee1 protein remains constant during G2 (Aligue et al., 1997; Keifenheim et al., 2017), suggesting that size-dependent mechanisms altering Wee1 activity and/or localization might exist. A recent study identified progressive phosphorylation of Wee1 as cells grow during G2, raising the possibility that inhibitory kinases might increasingly act on Wee1 as cells grow (Lucena et al., 2017).

Genetic and biochemical studies have identified two SAD family protein kinases, Cdr1 and Cdr2, which act as upstream inhibitors of Wee1. Both deletion and kinase-dead mutations in *cdr1* and *cdr2* result in elongated cells due to mis-regulation of Wee1 (Breeding et al., 1998; Kanoh and Russell, 1998; Russell and Nurse, 1987; Wu and Russell, 1993; Young and Fantes, 1987). Cdr1 can directly phosphorylate the Wee1 kinase domain to inhibit catalytic activity *in vitro* (Coleman et al., 1993; Parker et al., 1993; Wu and Russell, 1993). The role of Cdr2 kinase activity is less clear, but Cdr2 activation increases during cell growth in G2 (Deng et al., 2014). A key role for Cdr2 in this pathway is to assemble large, immobile “node” structures at the plasma membrane in the cell middle (Morrell et al., 2004). These interphase nodes are poorly defined oligomers of Cdr2, which then recruit Cdr1 to these sites (Guzman-Vendrell et al., 2015; Martin and Berthelot-Grosjean, 2009; Moseley et al., 2009). Wee1 primarily localizes in the nucleus and the spindle-pole body (SPB), where it can interact with Cdk1 (Masuda et al., 2011; Moseley et al., 2009; Wu et al., 1996). Wee1 has also been visualized at cortical nodes in some studies (Akamatsu et al., 2017; Moseley et al., 2009) but not in others (Masuda et al., 2011; Wu et al., 1996), and the low expression level of endogenous Wee1 has prevented careful analysis of its potential association with nodes.

Two models have been suggested to explain the relay of cell size information to Wee1 through Cdr2 nodes. The first model depends on the DYRK kinase Pom1, which directly phosphorylates Cdr2 to inhibit kinase activation and mitotic entry (Martin and Berthelot-Grosjean, 2009; Moseley et al., 2009). Pom1 forms size-invariant concentration gradients that emanate from the cell tips and could act as cellular “rulers,” but the concentration of Pom1 at the cell middle is largely independent of cell size, and*poml*Δ mutants retain size homeostasis (Bhatia et al., 2014; Hachet et al., 2011; Pan et al., 2014; Saunders et al., 2012; Wood and Nurse, 2013). An alternative model suggests that the accumulation of Cdr2 nodes, which double in number as cells double in size, generates size-dependent control of Wee1 (Pan et al., 2014). A limitation of previous studies has been the inability to monitor the signaling “output” of nodes because it has been unclear if, how, and when endogenous Wee1 localizes to nodes. Additionally, it has been unknown how nodes and their constituent proteins influence Wee1 phosphorylation *in vivo*.

Here, we found that nodes are stable, megadalton-sized complexes required for proper Wee1 phosphorylation in cells. By live-cell TIRF microscopy, we discovered that endogenous Wee1 localizes to Cdr2 nodes in transient bursts. The frequency and duration of bursts scale with cell size, and Pom1 regulates this node output specifically in small cells. Our combined results support a model whereby Cdr2 activity increases as cells grow, leading to size-dependent inhibition of Wee1 at nodes. This size-dependent pathway likely combines with other sizing mechanisms including Cdc25 accumulation to generate the homeostatic cell size control system.

## Results and Discussion

### Nodes are stable complexes that promote Wee1 phosphorylation

Cells require mechanisms to turn off Wee1 as they grow during G2 and approach the threshold size for division. Recent work identified progressive phosphorylation of Wee1 during this growth phase by monitoring a shift in Wee1 migration by SDS-PAGE (Lucena et al., 2017). Genetic studies place Cdr1 and Cdr2 kinases as upstream inhibitors of Wee1, but direct evidence of Wee1 phospho-regulation by Cdr1 and Cdr2 kinases in cells has been lacking. By monitoring the phosphorylation-dependent shift of endogenous Wee1 migration in western blots, we found that the most highly phosphorylated forms of Wee1 were absent in *cdr1*Δ mutants (Figure 1A). This result supports previous *in vitro* evidence for direct phosphorylation of Wee1 by Cdr1 (Coleman et al., 1993; Parker et al., 1993; Wu and Russell, 1993). Wee1 collapsed to a single band in *cdr2*Δ cells (Figure 1A), indicating a more dramatic loss of Wee1 phosphorylation in this mutant. These results show that Cdr1 and Cdr2 regulate phosphorylation of Wee1 in cells, and point to a key role for Cdr2 in this process. Cdr2 assembles into stationary nodes at the plasma membrane during G2, and recruits Cdr1 to these sites (Guzman-Vendrell et al., 2015; Martin and Berthelot-Grosjean, 2009; Morrell et al., 2004; Moseley et al., 2009). To test if assembly of Cdr2 and Cdr1 into nodes is important for Wee1 phosphorylation in cells, we blotted against Wee1 in *cdr2ΔC* cells. This mutant lacks the C-terminal KA1 domain that binds to both lipids and to Cdr1, but it retains the Cdr2 kinase and UBA domains that are required for catalytic activity. As a result, Cdr2ΔC does not bind the cortex or form nodes, and instead localizes diffusely to the cytoplasm and nucleus (Figure 1B) (Deng et al., 2014; Guzman-Vendrell et al., 2015; Moravcevic et al., 2010). We observed loss of Wee1 phosphorylation in the *cdr2ΔC* mutant similar to a *cdr2*Δ mutant (Figure 1C). We conclude that node-based signaling by Cdr1 and Cdr2 regulates phosphorylation of Wee1 in cells.

**Figure 1:**
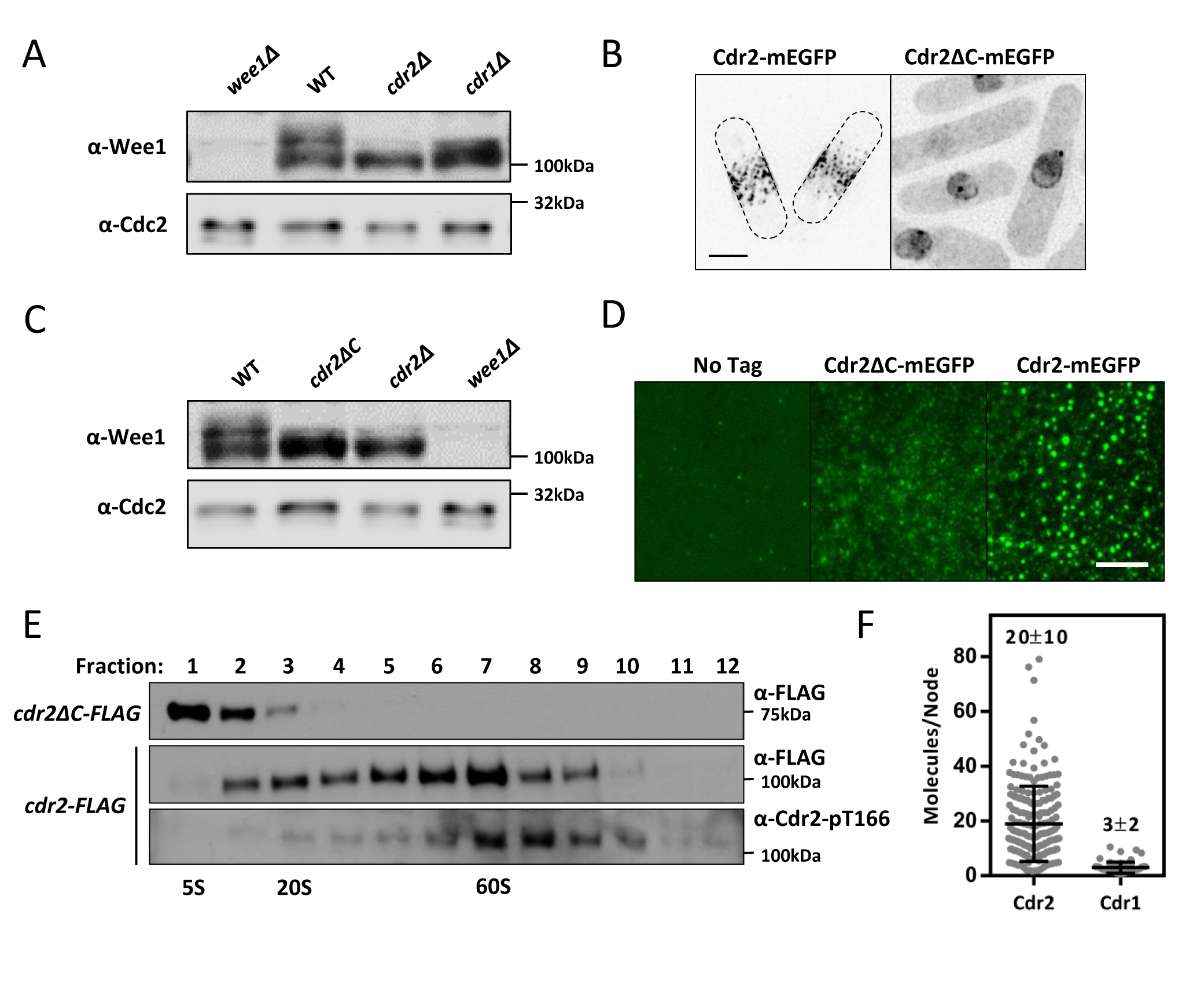
Nodes are stable scaffolds that facilitate inhibition of Wee1. (A) Wee1 phosphorylation is disrupted in *cdr1*Δ and *cdr2*Δ cells. Whole-cell extracts were separated by SDS-PAGE, and endogenous Wee1 was detected by western blot. (B) Localization of Cdr2-mEGFP and Cdr2ΔC-mEGFP in cells. Maximum intensity projections from Z-series are shown. Scale bar is 5μm. (C) Wee1 phosphorylation is disrupted in *cdr2ΔC* cells. Whole-cell extracts were analyzed as in panel A. (D) Visualization of node-like puncta by TIRF microscopy in whole-cell extracts from the indicated strains. Scale bar 5μm. (E) Detergent extracts from *cdr2-FLAG* or *cdr2ΔC-FLAG* cells were subjected to velocity sucrose gradient sedimentation, and fractions were probed against the FLAG tag or against Cdr2(pT166). Fraction 1 corresponds to the top of the gradient and contains smaller complexes; fraction 12 corresponds to bottom of the gradient. S-values were determined using size standards run on identical gradients. (F) Quantification of the number of Cdr2 and Cdr1 molecules per node (mean ± SD, n>65 each) based on super-resolution live-cell fluorescence microscopy.

The biophysical nature of Cdr2 nodes has been unclear. For example, these structures could represent stable biochemical complexes, or alternatively be comprised of weakly aggregated proteins. The critical role of nodes in Wee1 signaling led us to investigate their stability in cell lysates. Using TIRF microscopy, we observed node-like structures in the lysates of *cdr2-mEGFP* cells (Figure 1D). These structures were absent in the lysates of *cdr2ΔC-mEGFP* cells, and in lysates of control cells that do not express GFP (Figure 1D). This result suggests that nodes remain intact upon cell lysis and may represent stable complexes. To test this possibility, we performed sub-cellular fractionation of detergent extracts on velocity sucrose gradients. The mutant *cdr2ΔC* was found in early fractions representing monomeric proteins and/or small complexes (Figure 1E). In contrast, full-length Cdr2 migrated in a broad pattern with a peak at 60S (Figure 1E and S1A-B), consistent with a megadalton-sized complex. These fractions could be dialyzed and centrifuged on a second sucrose gradient without altering the sedimentation pattern of Cdr2 (Figure S1C), suggesting that nodes contain a stable core. TIRF microscopy of sucrose gradient fractions from Cdr2-mEGFP cells revealed node-like structures in later fractions, and diffuse signal in earlier fractions (Figure S1D). Thus, Cdr2 nodes represent large stable complexes as revealed by independent imaging and biochemical approaches.

Cdr2 is activated by phosphorylation of threonine 166 (T166) by the kinase Ssp1, and levels of Cdr2-pT166 increase with cell size (Deng et al., 2014). To determine the distribution of activated Cdr2 within our sucrose gradients, we used a phospho-specific antibody that detects Cdr2-pT166. We found that Cdr2-pT166 was enriched in the larger “node” fractions, and largely absent from the smaller fractions (Figure 1E). This suggests that nodes are sites for concentration of activated Cdr2. Large node fractions contain very low levels of Cdr1 (Figure S1E), suggesting it might not stably associate with the core of nodes. Consistent with this possibility, FRAP experiments revealed rapid turnover of Cdr1-mEGFP molecules at nodes *in vivo* (Figure S1F). We were unable to detect Wee1 in sucrose gradient experiments due to its low levels of expression and stability in lysates. These combined experiments identify a stable core of activated Cdr2 that forms cortical node structures for regulation of Wee1.

A second set of nodes, assembled by the protein Blt1, associate with Cdr2 nodes late in the cell cycle (Akamatsu et al., 2014; Moseley et al., 2009). Velocity sucrose gradients revealed the presence of Blt1 in a large protein complex in both wild type cells and *cdr2*Δ cells (Figure S1I). Similarly, Cdr2 still sedimented in node fractions in *blt1*Δ cells, which lack this second set of node structures (Figure S1I). This result is consistent with previous microscopy studies showing Cdr2 nodes in *blt1*Δ cells, and Blt1 nodes in *cdr2*Δ cells. Thus, Cdr2 forms a ~60S node complex independently of Blt1 nodes.

For comparison to our biochemical fractionation, we used quantitative super-resolution fluorescence microscopy to estimate the size of nodes in live cells. Using known measurements from previous studies (Wu and Pollard, 2005), we calibrated a confocal microscope to measure fluorescence intensity per mEGFP molecule (Figure S1G-H). With this set-up, we imaged *cdr2-mEGFP* and *cdr1-mEGFP* cells, and then calculated both global and local molecule numbers. The low expression level of Wee1-mEGFP prevented reliable *in vivo* molecule counting. Each cell contained an average of 5600 ± 1000 Cdr2-mEGFP molecules, and 700 ± 200 Cdr1-mEGFP molecules. More importantly, we used Airyscan confocal microscopy to measure 20 ±10 molecules of Cdr2 per node, and 3 ± 2 molecules of Cdr1 per node (Figure 1F), similar to independent studies using different imaging systems (Akamatsu et al., 2017; Pan et al., 2014). A node containing 20 Cdr2 would be nearly 2 megadaltons in size, and a globular complex of this size would sediment at approximately 60S, consistent with our sucrose gradient fractionation (Figure S1B). We note that both *in vitro* and *in vivo* approaches independently revealed a large spread in the size of nodes, which suggests that nodes can assume a range of sizes that could relate to their signaling properties. Overall, we conclude that nodes are large stable structures organized by an oligomeric core of Cdr2 molecules. These structures and their components are required for phosphorylation of Wee1 in cells.

### Wee1 localization to nodes requires Cdr2 kinase activity and the Wee1 N-terminus

To begin investigating how Wee1 is regulated by nodes, we tested its physical association with Cdr2. In a yeast two-hybrid assay, Wee1 was previously shown to interact with the Cdr2 kinase domain, and this interaction was lost in the catalytically inactive *cdr2(E177A)* mutant (Guzman-Vendrell et al., 2015). We confirmed this interaction, and then used it to determine the domain of Wee1 critical for Cdr2 interaction and for localization to nodes. The Wee1 protein is comprised of a C-terminal protein kinase domain and an N-terminal region of unknown function (Figure 2A). The N-terminal 150 residues of Wee1 were both necessary and sufficient to interact with Cdr2 in yeast two hybrid experiments, and these interactions were abolished by Cdr2(E177A) mutations (Figure 2B). This result is consistent with association of the Wee1 N-terminus with Cdr2 in cell lysates (Kanoh and Russell, 1998), and suggests the possibility that the Wee1 N-terminus functions to regulate localization to nodes. To test this idea, we examined localization of different Wee1 constructs after over-expression. Full-length Wee1 localized to nodes but Wee1(151-end), a truncated mutant lacking the Cdr2 association domain, did not localize to nodes (Figure 2C). Thus, the Wee1 N-terminus associates with Cdr2 and is required for localization to Cdr2 nodes. Although Wee1(1-150) was sufficient to interact with Cdr2 in the yeast two-hybrid assay, it was not sufficient for node localization in cells (Figure 2C), meaning that additional residues or structural features of Wee1 might contribute to its ability to localize to nodes upon over-expression.

**Figure 2:**
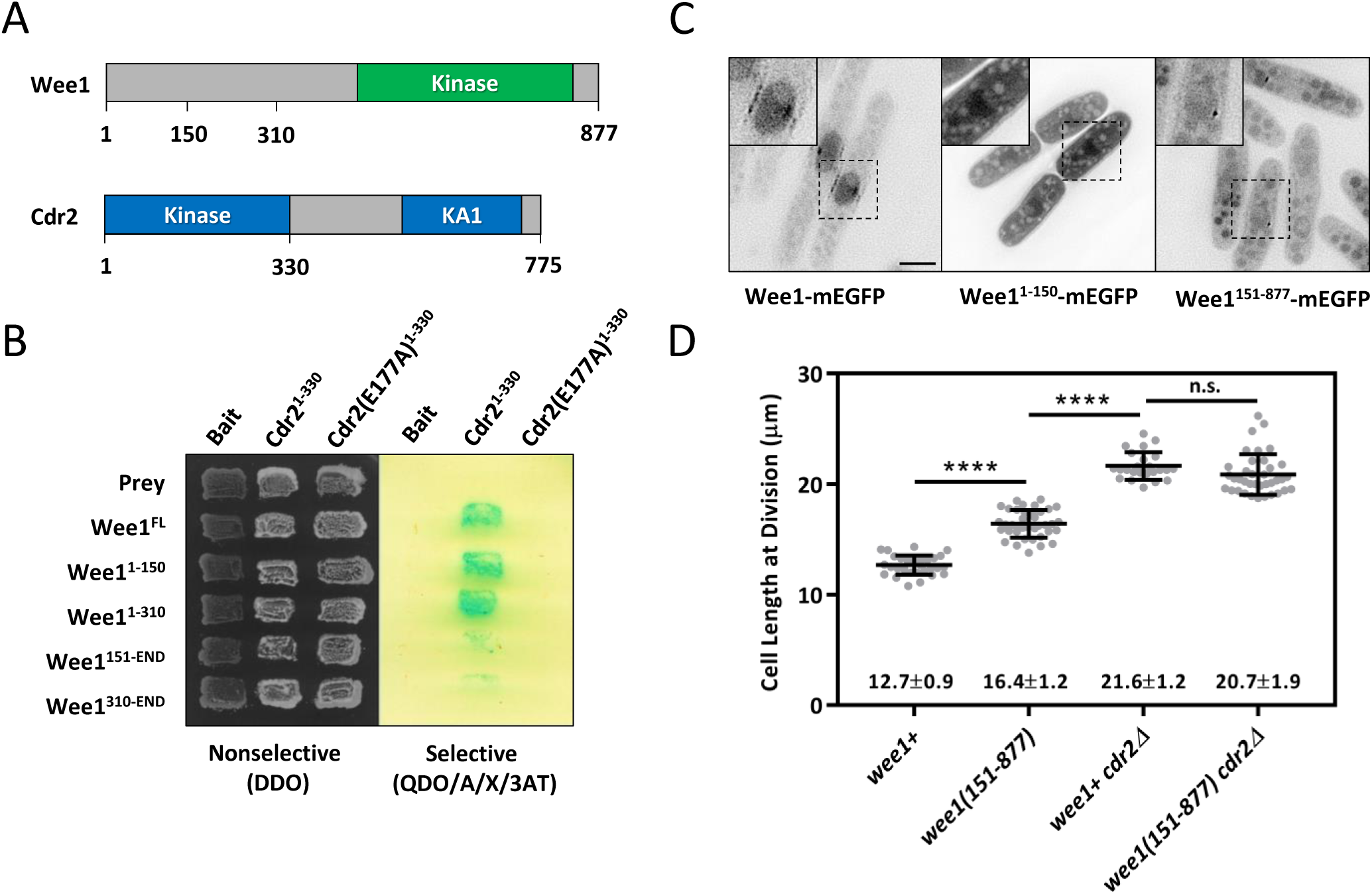
Wee1 localization to nodes requires Cdr2 kinase activity and the Wee1 N-terminus. (A) Schematic of Wee1 and Cdr2 functional domains. Values represent amino acid positions. (B) The Wee1 N-terminus interacts with wild type Cdr2 but not with kinase-dead Cdr2(E177A) in the yeast two-hybrid assay. Transformants were selected on a double dropout (DDO) plate, and interactions were tested on a quadruple drop-out plate containing aureobasidin, X-gal, and 3-AT (QDO/A/X/3AT) plate. Positive interactions are indicated by growth of blue colonies on selective plates. (C) Localization of the indicated Wee1 constructs over-expressed from the P81nmt1 promoter. Middle-focal plane wide-field images with inverted contrast are shown, and insets show enlarged view of dashed boxes. Scale bar 5μm. (D) Length of dividing, septated cells of the indicated genotypes (mean ± SD, n>50 cells). **** indicates *p* < 0.0001, n.s. indicates *p* > 0.05.

Next, we tested the functional role of the Wee1 N-terminus by integrating a truncation mutant expressed by the endogenous promoter as the sole copy of Wee1 in the cell. *wee1(151-end)* cells were elongated at division (Figure 2D), consistent with loss of Wee1 inhibition by the node pathway. This phenotype was not additive with *cdr2*Δ suggesting that the Wee1 N-terminus is required for regulation by Cdr2 (Figure 2D). In contrast, integrated full-length *wee1+* controls had no defects in cell size at division. These experiments show that the Wee1 N-terminus is required for interaction with Cdr2, localization to nodes, and negative regulation of Wee1 function.

### Dynamic bursts of Wee1 localization to Cdr2 nodes

Overexpressed Wee1 exhibits strong localization to Cdr2 nodes (Moseley et al., 2009), but localization of endogenously expressed Wee1 to nodes has been unclear. Wee1 is found in the nucleus and spindle pole body, where it can associate with its inhibitory target Cdk1 (Masuda et al., 2011; Moseley et al., 2009; Wu et al., 1996). Our results implicate nodes as sites of Wee1 regulation by Cdr2 and Cdr1, suggesting that endogenous Wee1 might localize to nodes. However, due to its low expression levels, there have been conflicting reports as to whether endogenous Wee1 localizes to cortical puncta (Akamatsu et al., 2017; Moseley et al., 2009) or does not localize to the cell cortex (Masuda et al., 2011). When seen at the cortex, low levels of Wee1 have prevented colocalization and time-lapse analyses. To study endogenous Wee1 at the cell cortex, we used the bright and photostable fluorophore mNeonGreen in combination with TIRF microscopy. In TIRF, we observed unambiguous Wee1-mNeonGreen localization at cortical puncta (Figure 3A). Strikingly, time-lapse imaging revealed that Wee1-mNeonGreen localized transiently to the cortex. From movies and kymographs, we found that Wee1 puncta bound and released the cell cortex with variable dwell times ranging from one frame (200ms) to over 1 minute (Figure 3A, Supplemental Video S1). Thus, endogenous Wee1 localizes to cortical nodes in transient bursts.

**Figure 3:**
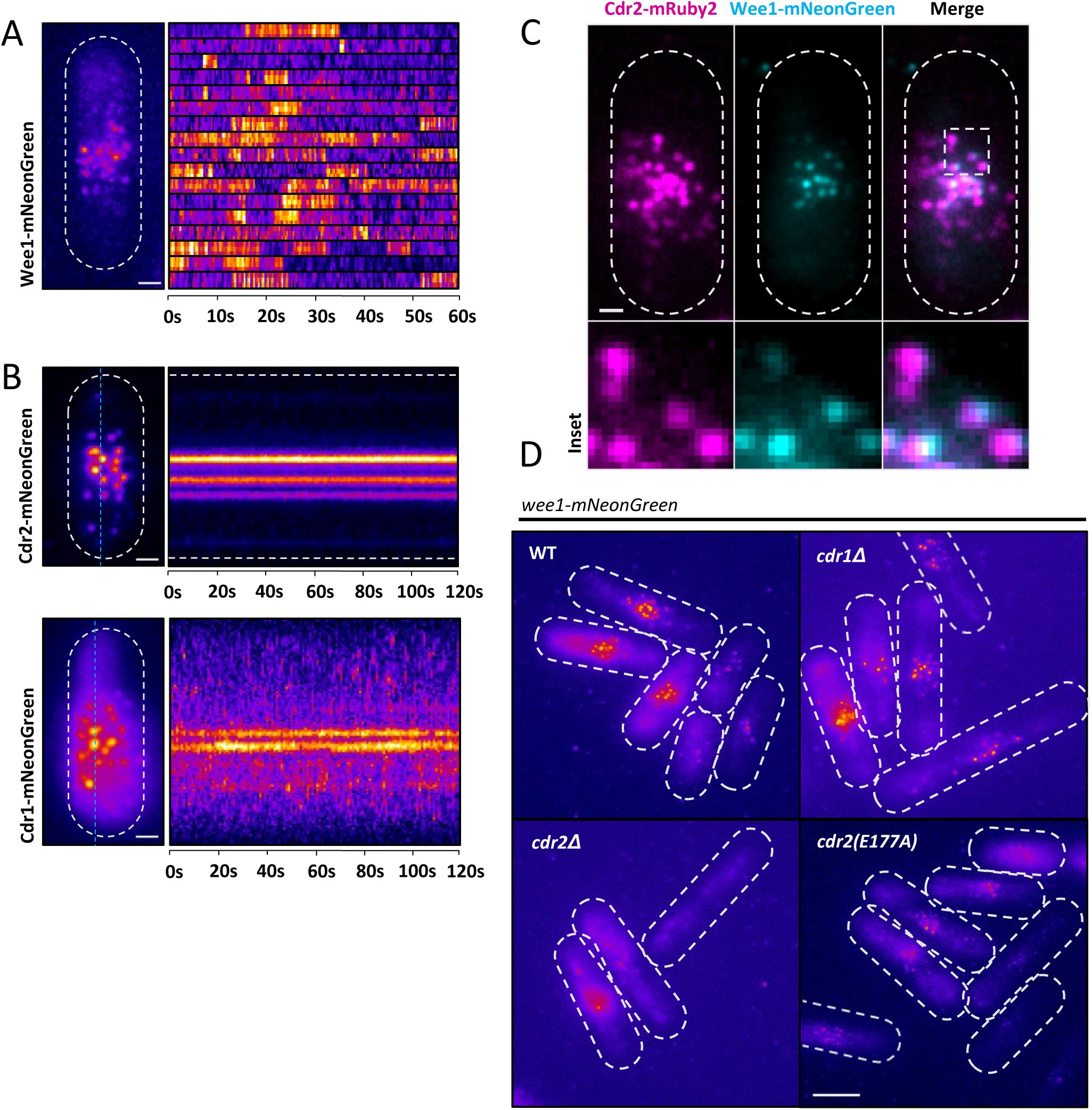
Wee1 puncta bind transiently to stable Cdr2 nodes. (A) Localization of Wee1-mNeonGreen by TIRF microscopy. Left panel is a maximum intensity projection of a 60s time-series imaged at 1s intervals. Scale bar is 1μm. Right panel is single-node 3x3 pixel kymographs of each Wee1 burst that appeared during the 60s time-lapse. See Supplemental Video S1. (B) Cdr2 and Cdr1 are stationary during 2m TIRF microscopy time-lapse acquisitions. Left panel images are maximum intensity projections of time-lapse acquisitions with 1s intervals. Right panels are kymographs of the time-lapse along the cyan-dashed line. Scale bar is 1μm. See Supplemental Videos S2 and S3. (C) Wee1 bursts colocalize with Cdr2 nodes. Images are dual-channel simultaneously-acquired TIRF microscopy images. Scale bar is 1μm. (D) Localization of Wee1 by TIRF microscopy in the indicated strains. Images are maximum intensity projections of 60s time-lapse, imaged at 1s intervals. Scale bar is 5μm.

The transient nature of Wee1 localization to cortical nodes in TIRF contrasts with other node proteins Cdr1 and Cdr2, which appear static in time-lapse movies from epifluorescence and confocal imaging systems. We performed four sets of experiments to support the conclusion that Wee1 transiently associates with stable Cdr2-Cdr1 nodes. First, we imaged endogenously tagged Cdr2-mNeonGreen or Cdr1-mNeonGreen in the same TIRF system. Both Cdr2-mNeonGreen and Cdr1-mNeonGreen persisted over long timescales at cortical nodes (Figure 3B, Supplemental Videos S2-S3), meaning that transient bursts are a specific behavior of wee1 at nodes and are not an artifact of TIRF imaging. Second, we imaged wee1 with a different fluorophore (wee1-mEGFP) and observed transient bursts, similar to wee1-mNeonGreen. Third, we tested if the position of the fluorophore at the Wee1 C-terminus induced transient bursts. We generated an N-terminal tag (mEGFP-wee1) and an in-frame internal tag (wee1-mEGFPint), integrated at the endogenous locus and expressed under control of the endogenous *wee1* promoter. These constructs also exhibited transient localization bursts to cortical nodes (Figure S2A-Bs). Fourth, using two-color TIRF we found that wee1-mNeonGreen bursts colocalized with Cdr2-mRuby2 nodes (Figure 3C) but not with other cortical structures like eisosomes or Skb1-Slf1 nodes (Figure S2D-E). These combined experiments confirm that wee1 localizes to Cdr2-Cdr1 cortical nodes in transient bursts, in contrast to other node components.

We next examined the molecular requirements for wee1 node bursts. wee1 puncta were unaffected in *cdr1*Δ cells but were absent from *cdr2*Δ cells (Figure 3D), which lack cortical nodes. wee1 puncta were also reduced in the kinase-dead *cdr2(E177A)* mutant (Figure 3D and S2I). We confirmed that these wee1 bursts were at Cdr2(E177A) nodes using two-color TIRFM on *wee1-mNeonGreen cdr2(E177A)-mKate2 cells* (Supplemental Figure S2C). Cdr2(E177A)-mEGFP still forms cortical nodes and still recruits Cdr1 to nodes (Morrell et al., 2004; Moseley et al., 2009). Further, Cdr2(E177A)-mEGFP nodes appeared identical to wild type by super-resolution molecule counting and FRAP experiments (Figure S2F-G). These results suggest that bursts of wee1 localization to inhibitory nodes might serve as a functional output for Cdr2-based kinase signaling. Further, regulation of Cdr2 kinase activity has the potential for dynamic control of wee1 through node bursts.

### The frequency of wee1 node bursts increases with cell size

Wee1 is progressively phosphorylated as cells increase in size during G2 phase (Lucena et al., 2017), and we have shown that phosphorylation requires the inhibitory kinases Cdr1 and Cdr2. If wee1 localization bursts to Cdr2 nodes mediate this progressive inhibitory phosphorylation, then we would predict size-dependent changes in wee1 node bursts. Consistent with this prediction, wee1 localization at nodes increased dramatically with cell size (Figure 4A). To quantify this increase, we measured the number of wee1-mNeonGreen puncta visible in TIRFM images, taken with a 1-second exposure, as a function of cell length. This analysis revealed a progressive increase in the number of wee1 nodes as a function of cell size, resulting in a 20-fold difference in wee1 nodes in large cells versus small cells (Figure 4B). Because many small cells lack detectable wee1 nodes and TIRF imaging is restricted to a single plane, this 20-fold increase may underestimate the increase over the entire cell surface.

**Figure 4:**
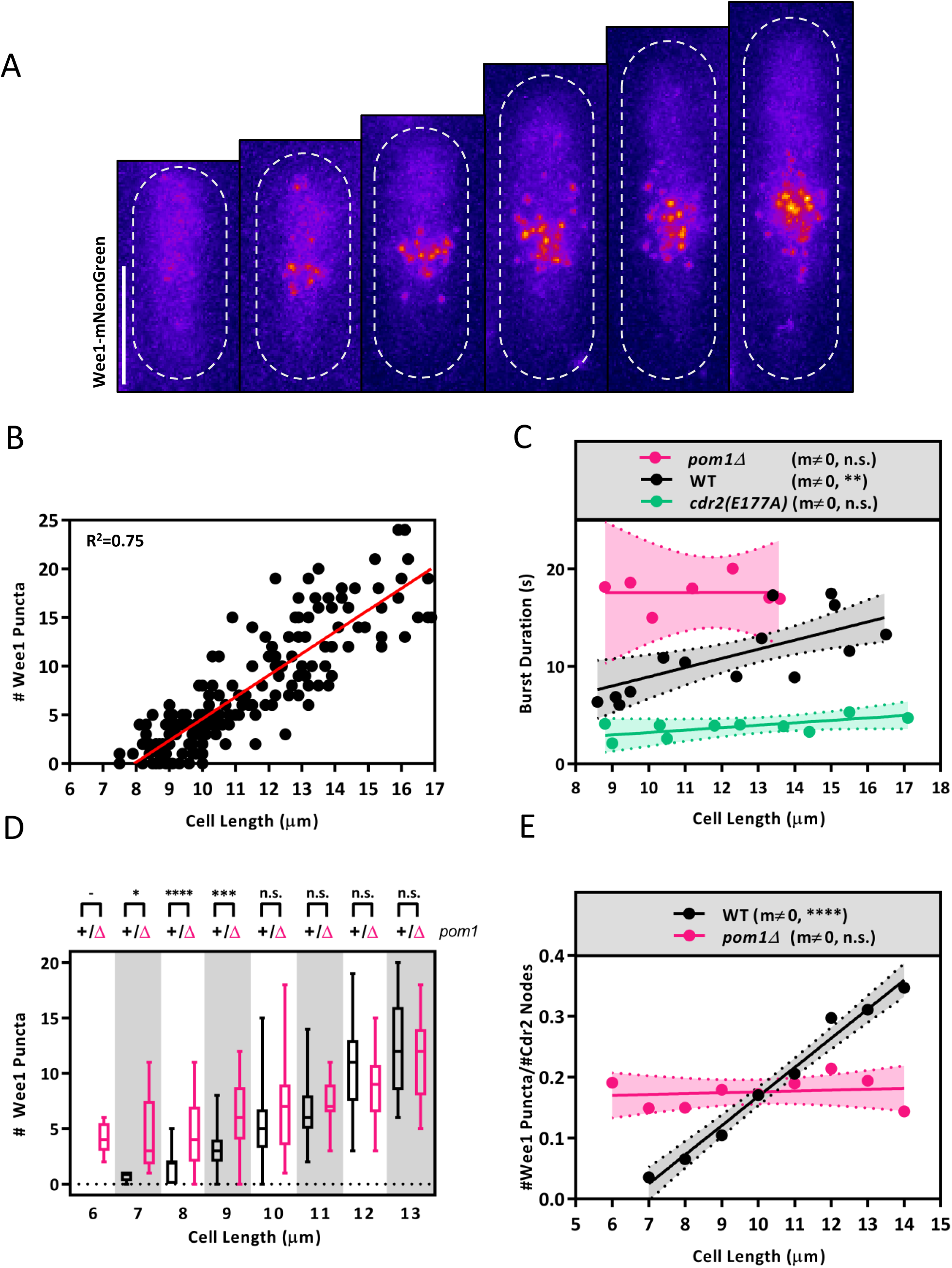
wee1 accumulation at nodes is size-dependent and buffered by Pom1. (A) Representative images of wee1 node localization in cells of increasing size imaged by TIRF microscopy. Scale bar is 5μm. (B) Quantification of the number of wee1 puncta as a function of cell size. Values were obtained from TIRF images with 1s exposure. Data fit a linear regression model. (C) wee1 burst duration scales with cell size in wild-type but not *cdr2(E177A)* or *pom1*Δ cells. The duration of individual wee1-mNeonGreen bursts was measured using time-lapse TIRF microscopy for cells of the indicated genotype. Dots correspond to mean burst duration plotted as a function of cell size. Lines are linear regressions fit to these data. Dashed lines indicate the 95% confidence interval of the linear regression, accounting for SD and N of the mean for each data point. The resulting slope is significantly non-zero (m ≠ 0) for wild type (*p* = 0.002, R^2^ = 0.02), but not for *pom1*Δ (*p* = 0.01, R^2^ = 0.01) or *cdr2(E177A)* (*p* = 0.9, R^2^ = 3.3e^-7^). (D) Pom1 suppresses wee1 bursts in small cells. The number of wee1 puncta was quantified as in panel B for WT and *pom1*Δ cells, and the data were binned according to cell size. Boxes indicate mean and SD, whiskers indicate min and max values. The number of wee1 bursts is significantly different between wildtype and *pom1*Δ in the three smallest bins (7μm: *p* = 0.01*, 8μm: *p* < 0.0001****, 9μm: *p* = 0.0003***, 10-13μm: *p* > 0.5). 6μm bin contains only *pom1*Δ cells. (E) Relative accumulation of wee1 puncta and Cdr2 nodes in WT and *pom1*Δ cells as a function of cell size. Linear regressions and 95% confidence intervals are shown. The slope of the linear regression is significantly non-zero for wildtype (*p* < 0.0001, R^2^ = 0.98) but not for *pom1*Δ (*p* = 0.7, R^2^ = 0.26).

Both the frequency and duration of wee1 bursts might be modulated to generate this cell size dependency. The number of Cdr2 nodes doubles during growth in G2 (Akamatsu et al., 2017; Deng and Moseley, 2013; Pan et al., 2014), which we corroborated by measuring the number of Cdr2-mEGFP nodes in the TIRF field as a function of cell size (Figure S3A). This two-fold increase in Cdr2 node number likely contributes to the 20-fold increase wee1 localization at nodes by increasing the frequency of total bursts within a cell. However, the difference in scale indicates that additional regulatory mechanisms must exist. Therefore, we asked if the duration of wee1 bursts changes as a function of cell size. Time-lapse TIRF microscopy of wee1-mNeonGreen revealed that the average dwell time of wee1 bursts is significantly correlated with cell size (slope of linear regression significantly non-zero, *p* = 0.002**). wee1 bursts in smaller cells lasted 6±6s (mean ± SD, n=4 cells in size range 7-9 μm), and this duration doubled to ~14±17s in large cells (Figure 4C, mean ± SD for 4 cells in size range 14-16 μm). Size-dependent accumulation of wee1 nodes was unaffected in *cdr1Δ* or *blt1Δ* cells, but was strongly reduced in kinase-dead *cdr2(E177A)* mutant cells (Figure S2H-K). Cdr2 kinase activity was also required for size-dependent change in wee1 burst duration. In *cdr2(E177A)* mutant cells, wee1 bursts lasted ~3.8s regardless of cell size (n=10, Figure 4C). This result indicates that Cdr2 kinase activity detains wee1 at a node. We conclude that increases in Cdr2 kinase activity and node number contribute to size-dependent regulation in wee1 bursts.

### Pom1 suppresses wee1 node bursts in small cells

We next tested how Pom1-Cdr2 signaling impacts size-dependent bursts of wee1 to nodes. Pom1 directly phosphorylates Cdr2 to inhibit mitotic entry through wee1. This modification inhibits node assembly and prevents the activating kinase Ssp1 from phosphorylating the Cdr2 activation loop (Bhatia et al., 2014; Deng et al., 2014; Guzman-Vendrell et al., 2015; Kettenbach et al., 2015; Moseley et al., 2009; Rincon et al., 2014). The levels of activated Cdr2 increase as cells grow, raising the possibility that Pom1 might inhibit Cdr2 activation and downstream signaling events in a size-dependent manner (Deng et al., 2014). In *pom1*Δ cells, the size dependence of wee1 node accumulation was partially disrupted due to increased wee1 localization at nodes in small cells (Figure 4D). wee1 puncta in large cells were unaffected by lack of Pom1. Because Pom1 inhibits Cdr2 kinase activity, which regulates the duration of wee1 bursts, we wondered if Pom1 acts in part by affecting the duration of wee1 bursts. In *pom1*Δ cells, wee1 burst duration was uniformly high (~17.7s) and was independent of cell size (Figure 4C, n=7). We conclude that Pom1 functions to reduce the duration of wee1 bursts to inhibitory Cdr2 nodes in small cells. This activity likely contributes to premature mitotic entry in *pom1* mutant cells, while additional mechanisms maintain size-dependent regulation of wee1 localization in larger cells.

Our data suggest that Pom1-dependent control of Cdr2 acts in combination with the 2-fold increase in Cdr2 node number, and these two combined factors generate the 20-fold increase in wee1 localization to nodes. To test this possibility, we measured how wee1 node bursts scale with Cdr2 node number in wild type versus *pom1*Δ cells. In wild type cells, the number of wee1 puncta scales more steeply with cell size than the number of Cdr2 nodes, suggesting Cdr2 node number alone does not explain the frequency of wee1 bursting. In contrast, in *pom1*Δ cells, the ratio of wee1 node bursts to Cdr2 node number was constant across of range of cell sizes (Figure 4E). Thus, in the absence of Pom1 regulation, the number of wee1 node bursts scales with the number of Cdr2 nodes: both double as cell size doubles. Pom1 acts within this context to inhibit Cdr2 kinase activity and thereby suppress wee1 node bursts in small cells. The effect of this Pom1 activity is to magnify the fold increase in wee1 inhibition as cells double in size. These combined mechanisms contribute to the ~20-fold increase in wee1 bursts to inhibitory nodes as cells double in size.

Our work presents a framework to understand the signaling steps that occur within a Cdr2 cortical node (Figure 5). We found that Cdr2 kinase activity regulates wee1 localization to nodes, while previous work demonstrated that Cdr1 inhibits wee1 kinase activity through phosphorylation (Coleman et al., 1993; Parker et al., 1993; Wu and Russell, 1993). These distinct regulatory mechanisms act on different domains of the wee1 protein. Cdr2 phosphorylates the N-terminal domain that we found is critical for node localization (Kanoh and Russell, 1998), while Cdr1 phosphorylates the C-terminal kinase domain of wee1 (Coleman et al., 1993; Parker et al., 1993; Wu and Russell, 1993). These results raise the possibility that initial phosphorylation by Cdr2 traps wee1 molecules at a node for several seconds, allowing subsequent phosphorylation by Cdr1, which results in a catalytically inactive wee1 molecule. Such processive phosphorylation would be consistent with partial loss of wee1 phosphorylation in *cdr1*Δ cells, with more severe loss of phosphorylation in *cdr2*Δ cells. Future work will be directed at mapping residues phosphorylated by Cdr1 versus Cdr2, and elucidating the ultrastructural organization of these molecules within megadalton-sized nodes. We also note that nodes represent sites of wee1 inhibition, while the nucleus and SPB are sites of wee1 overlap with Cdk1. These two pools likely exchange through nuclear shuttling, which presents another potential point of regulation as cells grow and progressively inhibit wee1. As cells grow, the increased frequency of wee1 bursts to inhibitory nodes likely decreases cellular wee1 activity, thus contributing to Cdk1 activation for mitotic entry.

**Figure 5:**
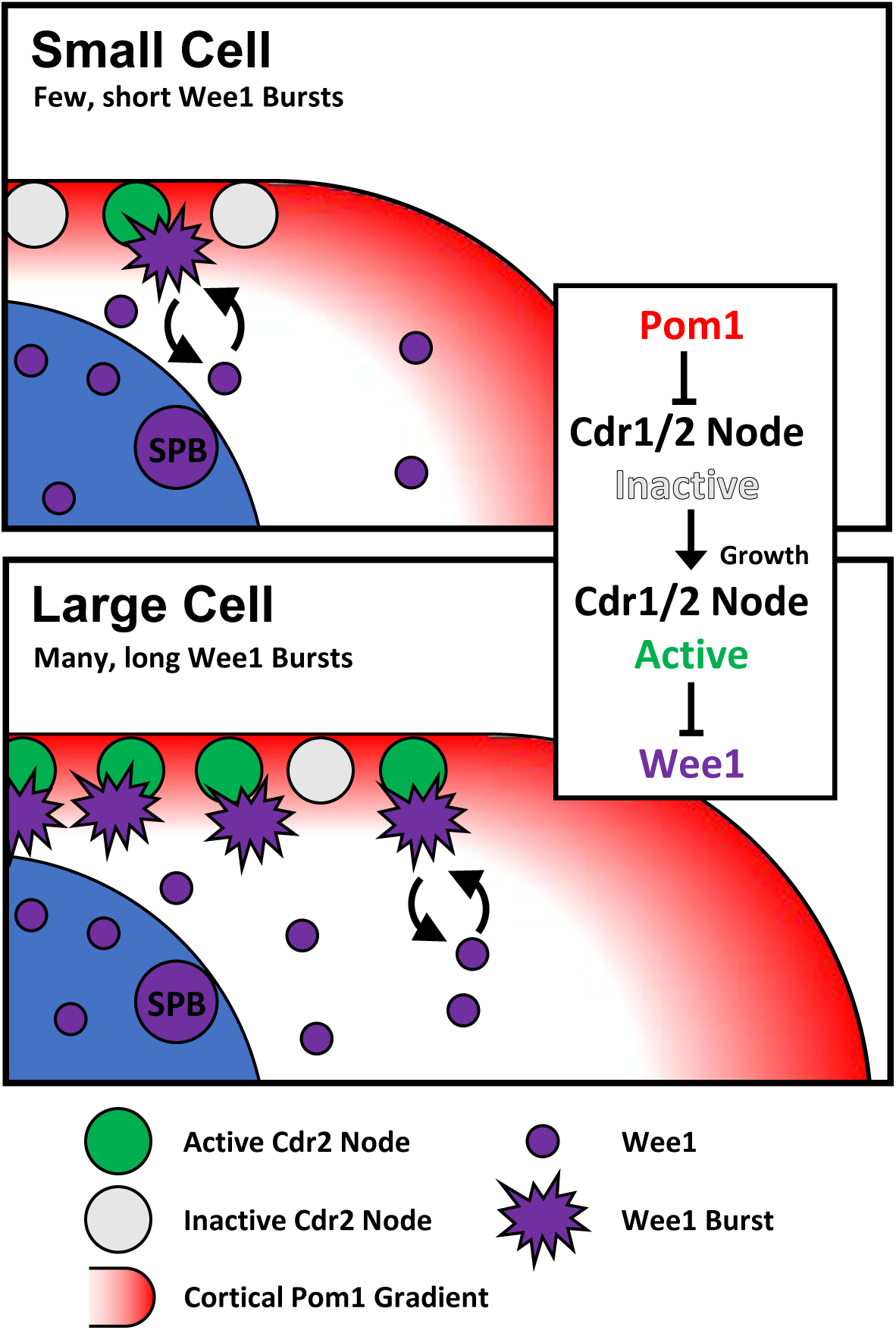
A working model for size-dependent control of wee1 by bursts of node localization. See the text for discussion.

Our data support a model whereby wee1 regulation by the Cdr2-Cdr1 pathway is cell size-dependent due to the combination of Pom1 activity in small cells and Cdr2 node accumulation. Mutations in this pathway alter cell size at division but do not impair cell size homeostasis (Wood and Nurse, 2013). This is not surprising given the existence of other size-dependent signaling mechanisms, such as accumulation of the mitotic inducer Cdc25 (Keifenheim et al., 2017; Moreno et al., 1990), which also may not be solely responsible for cell size homeostasis. We consider it likely that cell size homeostasis is a robust, systems-level property that emerges from the integration of multiple size-dependent signaling pathways, including the node-based mechanism studied here. In this sense, it will be interesting to uncover how these different mechanisms connect, converge, and alter each other’s signaling properties. Finally, cell size is modulated by nutrient availability in fission yeast and other organisms. The *cdr2* and *cdr1* mutants were initially identified due to altered cell cycle regulation upon nitrogen stress (Feilotter et al., 1991; Young and Fantes, 1987), and the Pom1-Cdr2 pathway was recently implicated in cell cycle control during glucose deprivation (Kelkar and Martin, 2015). How environmental signals control wee1 node bursts or other aspects of node-based signaling may reveal new mechanisms within this network.

## Materials and Methods

### Strain Construction and Media

Standard *S. pombe* media and methods were used (Moreno et al., 1991). Strains used in this study are listed in Supplementary Table S1. For cell length measurements, >50 septated cells from log-phase cultures grown in YE4S at 25C were measured by Blankophor staining. Gene tagging and deletion were performed using PCR and homologous recombination (Bähler et al., 1998). Truncation of the endogenous wee1 N-terminus was performed by first deleting AA1-150 from plasmid pJK148-Pwee1-wee1-Twee1 using QuikChange II mutagenesis (Stratagene). The parental and mutated plasmids were linearized by NruI digestion and integrated into the leu1 locus of diploid strain JM712 *wee1+/wee1Δ::kanMX6 leu1-32/leu1-32 ura4-D18/ura4-D18 ade6- M210/ade6-M210 h+/h+* and selected on EMM-Leu plates. The resultant diploid strain was transformed with the sporulation-inducing plasmid pON177 (h- mating plasmid with ura4 selectable marker - pJM363), and spores were separated by tetrad dissection and selected on YE4S+G418 and EMM-Leu plates. The mNeonGreen sequence was used under license from Allele Biotechnology. We concluded that this tag did not impair wee1 function because *wee1-mNeonGreen* cells divided at 14.6± 1.1 μm in length (mean ± SD for n>50 cells grown in EMM4S at 32°C).

### Sucrose Gradient Ultracentrifugation

Fission yeast detergent extracts were prepared by growing 1.5L cells to mid-log phase (OD ~0.3), and washing 2X with 50mL Node Isolation Buffer (NIB – 50mM HEPES, 100mM KCl, 1mM MgCl2, 1mM EDTA, pH 7.5). The pellet was then resuspended in an equal volume of 2X NIB (W/V) containing a protease/phosphatase inhibitor cocktail (10μL/mL 200x PI, 50μL/mL 1M β-glycerol phosphate, 50μL/mL 1M NaF, 2.5 μL/mL 200mM PMSF, 1μL/mL 1M DTT) and snap frozen as pellets by pipetting into a liquid nitrogen bath. Yeast pellets were then ground using liquid-nitrogen chilled coffee-grinders for 2 min, and collected into chilled falcon tubes and stored at −80°C. 1.5g of frozen yeast powder was then thawed on ice, and Triton X-100 was added to a final concentration of 1%, and the extracts were mixed by gentle pipetting. Extracts were then centrifuged at 4^0^C for 10min at 20,000 x g to yield a low speed supernatant, which was then subjected to sucrose gradient fractionation.

Discontinuous sucrose gradients were prepared by layering 5 – 23% (top to bottom) sucrose in NIB + 1% Triton X-100 in 1.2mL steps of 2% sucrose increments. 400μL low-speed detergent extracted supernatant was then added to the top of the gradient. Sucrose gradients were centrifuged at 100k x g (35kRPM) in a Beckman L8-M ultracentrifuge for 3.5hr at 4°C in a chilled SW41 swinging bucket rotor. 1mL gradient fractions were collected from the top by hand, vortexed, and 100μL of each fraction was mixed 2:1 in 3X western blot sample buffer (65 mM Tris pH 6.8, 3% SDS, 10% glycerol, 10% 2-mercaptoethanol, 50 mM NaF, 50 mM β-glycerophosphate, 1 mM sodium orthovanadate) and boiled for 5 min. Samples were then subjected to SDS-PAGE and western blotting.

### Western Blotting

For gel-shift western blots, whole cell extracts were lysed in 100 μl of 3X western blot sample buffer in a Mini-beadbeater-16 (Biospec) for 2 minutes. Gels were run at a constant 20 mAmps until 75kDa marker was at the bottom of the gel. Blots were probed with anti-FLAG M2 (Sigma) and anti-wee1 (Deng and Moseley, 2013). For wee1 western blots, we also repeated each experiment with a different anti-wee1 antibody (Lucena et al., 2017), which was provided by Douglas Kellogg and yielded identical results (not shown). For monitoring wee1 phosphorylation, samples were run on an SDS-PAGE a gel containing 6% acrylamide and 0.02% bisacrylamide.

For sucrose-gradient western blots, samples were prepared as described and run at constant-voltage of 100mV on 10% acrylamide gels. Blots were probed with anti-FLAG M2 or with anti-Cdr2-pT166 (Deng et al., 2014). To reduce background signal, nitrocellulose membranes for phospho-specific western-blots were incubated in 4% BSA/TBST for both primary and secondary antibodies at 4°C, with 2 hour washes in between.

### Yeast Two-Hybrid

A Matchmaker yeast two-hybrid system (Clontech) was used to test for physical interactions between Cdr2 and wee1 constructs. Bait plasmids were generated by ligating PCR amplified fragments into pGBKT7 using XmaI/SalI restriction sites. Prey plasmids were generated by ligating PCR fragments into pGADT7 using NdeI/XmaI restriction sites. All bait and prey plasmids were transformed into budding yeast strain Y2H-Gold on single-drop out (SDO) plates (SD-Leu or Trp), and tested individually for autoactivation of the reporters on quadruple-drop out plates (SD-Leu-Trp-Ade-His) with X-gal, 125ng/mL aureobasidin and 30mM 3-amino-1,2,4-triazole (QDO/X/A/3AT). The full length Cdr2 bait construct induced autoactivation, and so we used a truncated construct containing Cdr2 AA1-330 to test pair-wise interactions. For interaction tests, bait and prey plasmids were co-transformed into Y2H-Gold cells and selected on DDO plates (SD-Trp-Leu) before scoring of interactions on QDO/X/A/3AT plates.

### Widefield Microscopy and Analysis

Cells in Figure 1B were imaged in liquid EMM4S using a DeltaVision Imaging System (Applied Precision) composed of an Olympus IX-71 inverted wide-field microscope (Olympus), a CoolSNAP HQ2 camera (Photometrics), and an Insight solid-state illumination unit (Applied Precision). Z-stacks were acquired at 0.5μm intervals and processed by iterative deconvolution using SoftWoRx software (Applied Precision). Two-dimensional maximum intensity projections of these Z-stacks were rendered using ImageJ (National Institutes of Health).

### TIRF Microscopy and Analysis

Node components were imaged using simultaneous dual-color total internal reflection fluorescence microscopy to limit excitation of fluorophores to those nearest to coverslip. Imaging was performed on a commercially available TIRF microscope from Micro Video Instruments (MVI) equipped with a 100x Nikon Apo TIRF NA 1.49 objective and a two-camera imaging adaptor (Tu-CAM, Andor Technlogy) containing a dichroic and polarization filters (Semrock FF580-FDi01-25x36, FF02-525/40-25, FF01-640/40-25) to split red and green signal between two aligned Andor iXon electron-multiplied CCD cameras (Andor Technology). Red/green beam alignment was performed prior to imaging using a TetraSpeck Fluorescent Microsphere size kit (Thermofischer).

Standard #1.5 glass coverslips were RCA cleaned before use to remove fluorescent debris. Cells were grown in EMM4S, and washed into fresh EMM4S immediately before imaging to remove auto-fluorescent puncta resulting from overnight culture. Cells were imaged in EMM4S media on glass slides. Individual slides were used for no more than five minutes to prevent cells from exhausting nutrients or oxygen. Agar pads were not used due to increased background fluorescence.

Image analysis and processing was performed using ImageJ (NIH). Cdr2 node numbers and wee1 puncta number and binding duration were quantified using the MosaicSuite ParticleTracker plugin (Sbalzarini and Koumoutsakos, 2005). Due to variable fluorescence intensity in different TIRF fields and images, thresholding parameters were determined separately for each image, and accuracy was confirmed by visual inspection to ensure that only nodes/puncta were counted and that no nodes/puncta were omitted. Particle diameter was set to 1 pixel. Linking range for particle tracking was set to 2 pixels, because neither Cdr2 nodes nor wee1 puncta exhibit appreciable cortical diffusion on the time scale of our imaging. Lookup table was adjusted to fire for images in Figures 3, 4, and S2 to emphasize signal intensities. Statistical significance of differences in wee1 burst accumulation (Figure 4D) was determined using a Welch’s T-test.

### Single-molecule counting

Local and cellular molecule counting were performed as in (Wu and Pollard, 2005). To determine the number of Cdr1 or Cdr2 molecules per node, we calibrated a fluorescence microscope to determine the local number of mEGFP molecules based on fluorescence intensity. A standard curve relating fluorescence intensity to the number of molecules per structure was created by imaging mEGFP-tagged Cdc12, Spn1, or Sad1, which have known numbers of molecules at their respective structures. These strains were provided by Thomas Pollard. We then imaged Cdr2-mEGFP or Cdr1-mEGFP in the same genetic background. Cells were grown in YE4S to mid-log phase, concentrated, and imaged on YE4S agar pads using identical imaging parameters. To achieve maximum resolution and sensitivity, imaging was performed on a commercially available Zeiss LSM-880 laser scanning confocal microscope equipped with Airyscan super-resolution module and GaAsP Detectors, using the resolution vs. sensitivity mode to optimize signal to noise. For each strain, we collected 40 slice Z-stacks with 0.17μm spacing to image completely through entire cells with high spatial resolution in all dimensions. Airyscan images were processed in Zeiss Zen Blue software, and quantification was performed on sum projections of Airyscan reconstructed stacks. For preparation of the standard curve, integrated fluorescence density of Cdc12-meGFP at cytokinesis nodes or contractile rings, Sad1-mEGFP at spindle pole bodies, and Spn1-mEGFP at septin rings was measured, and background signal was subtracted using an equal sized ROI in ImageJ. Fluorescence values were then plotted against their known molecular counts to generate the standard curve. Fluorescence intensity of Cdr2- mEGFP or Cdr1-mEGFP at single node ROIs was measured using identical imaging conditions and analysis, and their fluorescence values were plotted against the best-fit line of the standard curve to infer the number of molecules per node.

For determination of the cellular concentrations of Cdr1 and Cdr2, we similarly calibrated a Quorum Wave FX-X1 spinning disk confocal system (Quorum Technologies) and Hamamatsu ImageEM camera. A standard curve relating fluorescence intensity to cellular concentration was created by imaging mEGFP-tagged Cdc12, Spn1, Sad1, Ain1, Rlc1, or Mid1, which have known cellular concentrations. These strains were provided by Jian-Qiu Wu. We then imaged Cdr2-mEGFP or Cdr1-mEGFP in the same genetic background. Cells were grown in YE4S to mid-log phase, concentrated, and imaged on YE4S agar pads using identical imaging parameters. We collected a Z-stack comprised of 13 slices with 0.5μm spacing with the shutter open between steps. Sum projections were created from the entire stack, corrected for uneven illumination and processed to remove background and auto-fluorescence. Fluorescence intensity was measured from whole cell ROIs using ImageJ. Fluorescence values were then plotted against their known molecular counts to generate the standard curve. Fluorescence intensity of Cdr2-mEGFP or Cdr1- mEGFP cells was measured using identical imaging conditions and analysis, and their fluorescence values were plotted against the best-fit line of the standard curve to infer the number of molecules per cell.

### Fluorescence Recovery After Photobleaching (FRAP)

FRAP analysis was performed using a Zeiss LSM-880 (See Single Molecule Counting) using the Resolution vs. Sensitivity mode to decrease photobleaching during time-lapse and to optimize signal to noise. Cells were grown in YE4S to mid-log phase, concentrated, and imaged on YE4S agar pads. At each time interval, three optical-section Z-stacks with 0.16μm spacing for 0.48μm total thickness were acquired, processed with linear 2-dimensional Airyscan reconstruction, and used to generate sum-projection images of each time point for quantification using ImageJ. Bleaching was performed using the Zen Black FRAP software to bleach ROIs drawn around the node containing region on one side of the cell. Unbleached cells were used to correct for photobleaching of the sample during acquisition. Data were fit to one-phase association curves from which the mobile fraction and T_1/2_ were calculated.

---

**Supplemental Figure S1:**
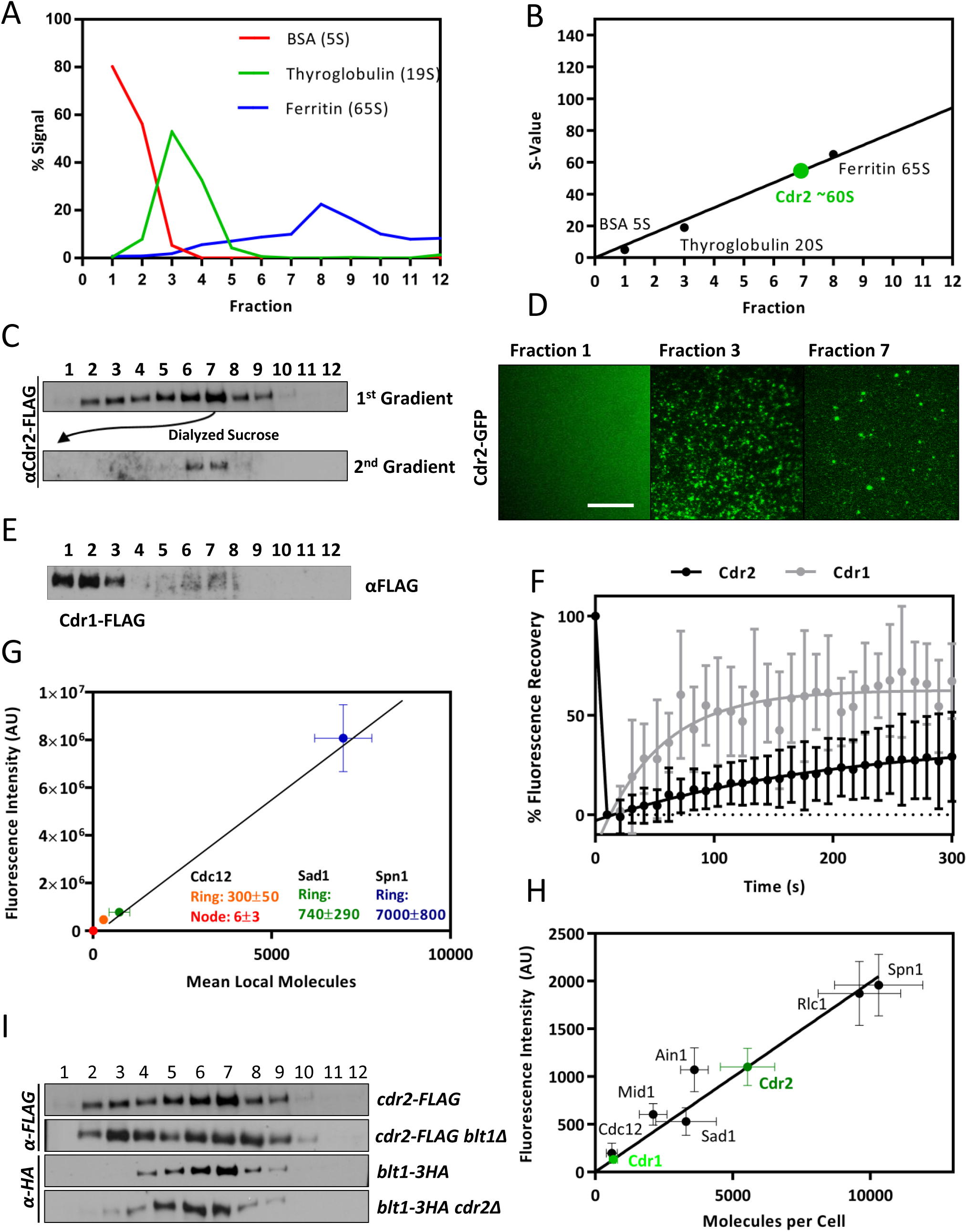
(A) Size standards for velocity sucrose gradients, as in main Figure 1E. (B) Plot of S-value versus sedimentation pattern. Line is linear regression of the three size standards, with the S-value for Cdr2 interpolated. (C) Node complexes remain stable through multiple sucrose gradients. Fraction 7 was dialyzed to remove sucrose, and then run on a second identical gradient with no change to sedimentation pattern. Blots were probed with anti-FLAG antibodies. (D) TIRF microscopy of sucrose gradient fractions from *cdr2-mEGFP* detergent extracts. Scale bar 5μm. (E) Velocity sucrose gradient sedimentation pattern of extracts prepared from *cdr1-5xFLAG* cells. Blots were probed with anti-FLAG antibodies. (F) Fluorescence recovery after photobleaching at cortical nodes for *cdr2-mEGFP* (n=17 cells) or *cdr1-mEGFP* (n=19 cells) cells. Data points are means, error bars show SD. Cdr2 t_1/2_ = 139s, Mobile Fraction = 0.38. Cdr1 t_1/2_ = 37s, Mobile Fraction = 0.63. (G) Standard curve for local single-molecule counting showing mean fluorescence intensity (A.U.) and mean number of molecules of the indicated protein/structure. Error bars indicate S.D. Linear regression used to interpolate experimental values is shown. Values used for standards are from (Wu and Pollard, 2005). (H) Standard curve for whole-cell molecule counting. Error bars indicate S.D. Linear regression used to interpolate experimental values is shown. Values used for standards are from (Wu and Pollard, 2005). (I) Western blots against Cdr2 and Blt1 node proteins in sucrose-gradient fractions prepared from the indicated strains.

**Supplemental Figure S2:**
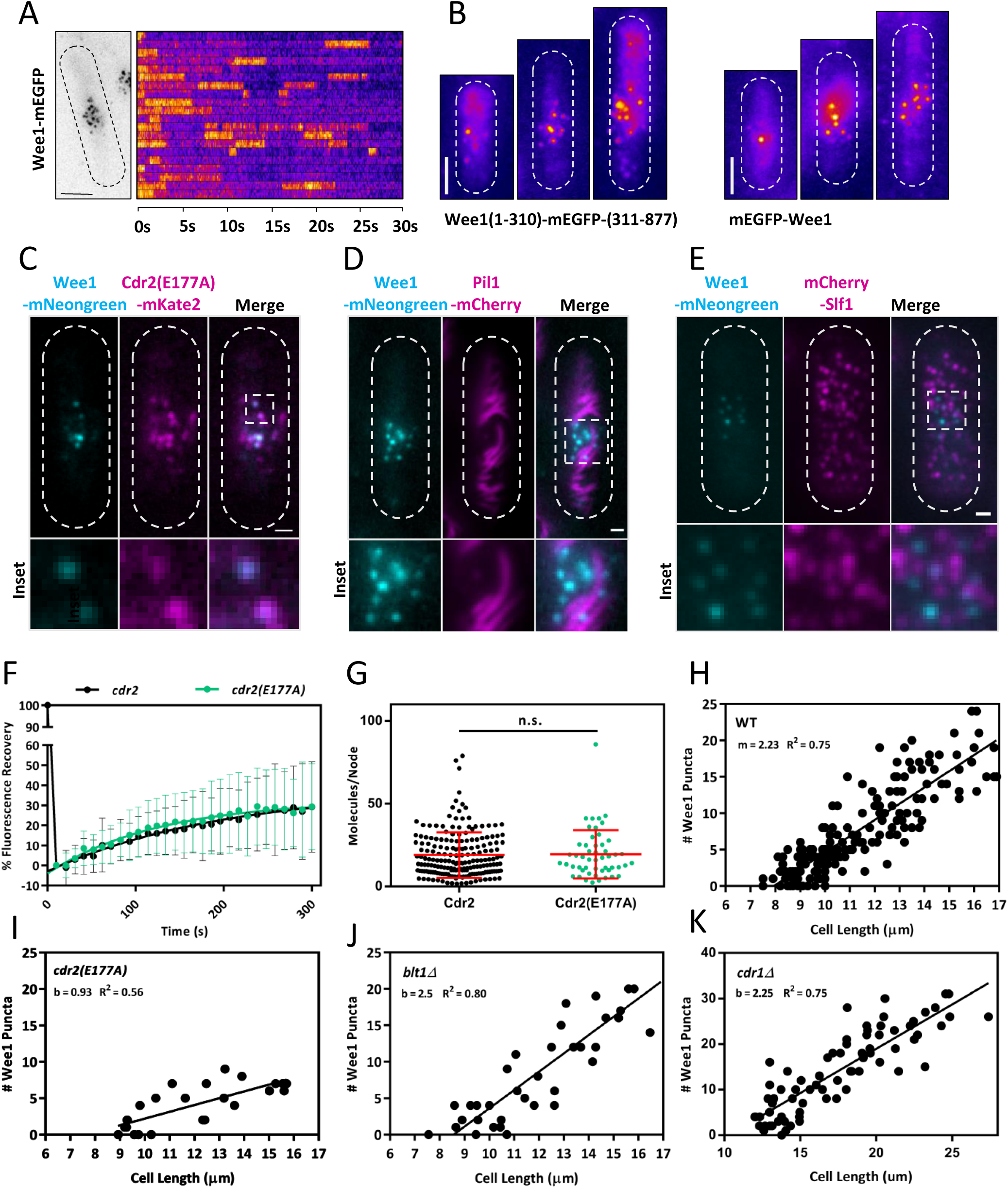
(A) Localization of wee1-mEGFP by TIRF microscopy. Left panel is a maximum intensity projection of 30s time-lapse for the outlined cell. Right panel is a compilation of 3x3 pixel single node kymographs for each wee1 burst in the acquisition. Scale bar is 4μm. (B) The location of the mEGFP tag does not affect localization of wee1 to nodes, as visualized by TIRF microscopy. Scale bar is 4μm. Cells in top row have internal mEGFP tag, cells in bottom row have N-terminal mEGFP tag. (C) wee1-mNeonGreen puncta colocalize with kinase-dead Cdr2(E177A)-mKate2 nodes by TIRF microscopy. Scale bar is 1μm. Lower boxes are zoom-images of the indicated region. (D-E) wee1-mNeonGreen puncta do not colocalize with the eisosome protein Pil1 or Slf1 nodes by TIRF microscopy. Scale bar is 1 μm. Lower boxes are zoom-images of the indicated region. (F) FRAP curves are unchanged in *cdr2-mEGFP* versus kinase-dead *cdr2(E177A)-mEGFP* cells. *cdr2-mEGFP* t_1/2_ = 139s, Mobile Fraction = 0.38; *cdr2(E177A)-mEGFP* t_1/2_ = 87s, Mobile Fraction = 0.32. Values not significantly different at 95% confidence level. Cdr2 data are the same as Figure S1. (G) The number of molecules of Cdr2 per node is unchanged between wild-type and *cdr2(E177A)* cells (n.s.,*p* = 0.83). Data for Cdr2 are the same as in Figure 1F, and replotted for comparison. (H) Size-dependent accumulation of wee1 puncta in WT cells. Data are replotted from Figure 4B for comparison to panels I-K. (I-K) Size-dependent accumulation of wee1 punta in *cdr2(E177A)*, *blt1*Δ, and *cdr1*Δ cells. Lines are linear regressions fit to the data.

**Supplemental Figure S3:**
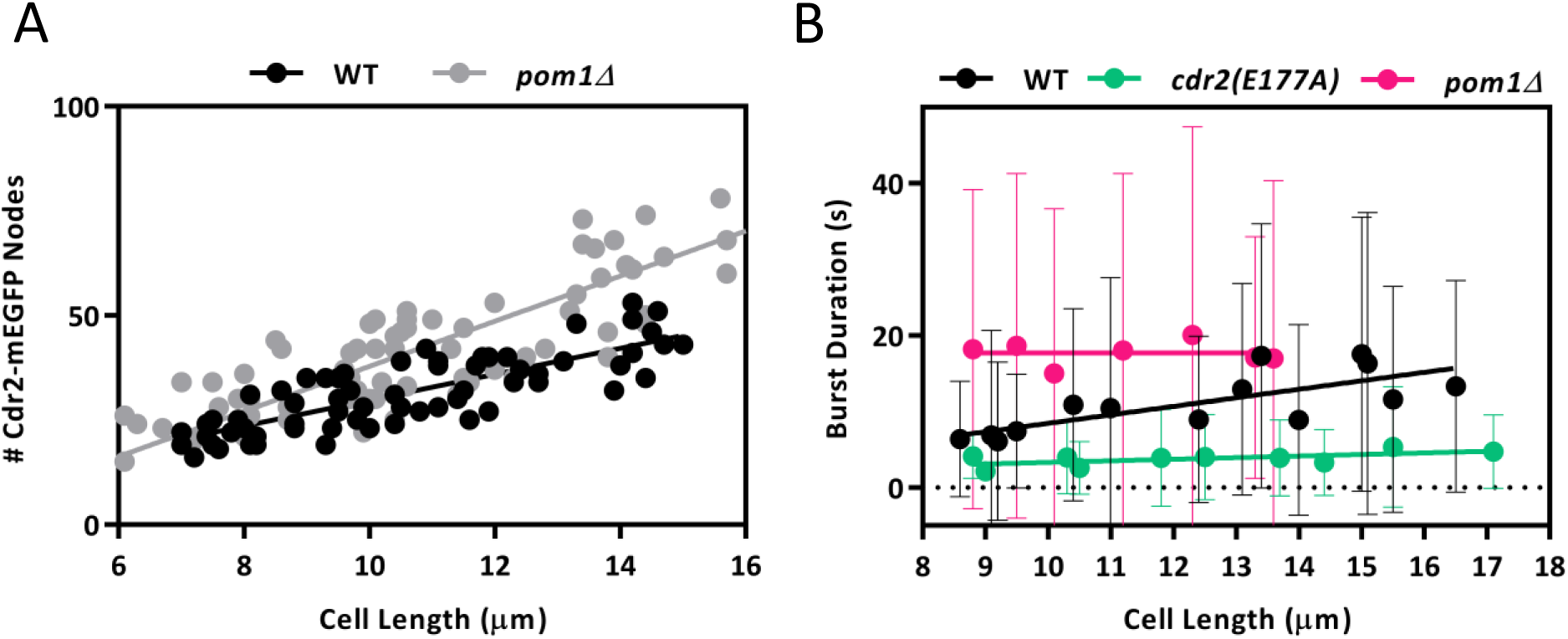
(A) Size-dependent accumulation of Cdr2-mEGFP nodes in wild-type versus *pom1Δ* cells. The number of Cdr2 nodes doubles with size in wild-type cells, but approximately triples in *pom1*Δ cells. The number of nodes was counted from TIRF images and therefore does not represent whole-cell counts. Lines are linear regressions of the data (WT R^2^ = 0.65, *pom1*Δ R^2^ = 0.74). (B) wee1 burst duration scales with cell size in wild type but not *cdr2(E177A)* or*pom1*Δ cells. The duration of individual wee1 bursts was measured using time-lapse TIRF microscopy of cells expressing wee1-mNeonGreen at endogenous levels in cells of the indicated genotype. Dots correspond to mean burst and S.D. of burst duration plotted as a function of cell size. Lines are linear regressions fit to these data. Data are the same as Figure 4C, and are re-plotted to present S.D. of the mean.

Supplemental Table S1: Fission yeast strains used in this study.

Supplemental Video S1: Timelapse TIRFM of the *wee1-mNeonGreen* cell shown in Figure 3A. 200 msec exposures were obtained at 1-sec intervals for 60 seconds. Scale bar is 1 μm.

Supplemental Video S2: Timelapse TIRFM of the *cdr2-mNeonGreen* cell shown in Figure 3B. 200 msec exposures were obtained at 1-sec intervals for 120 seconds. Scale bar is 1 μm.

Supplemental Video S3: Timelapse TIRFM of the *cdr1-mNeonGreen* cell shown in Figure 3B. 200 msec exposures were obtained at 1-sec intervals for 120 seconds. Scale bar is 1 μm.

## Acknowledgements

We thank members of the Moseley lab for comments on the manuscript, as well as B. Wickner and C. Barlowe at Dartmouth for sharing equipment. We thank members of the A. Gladfelter lab and A. Lavanway for assistance with microscopy; A. Orr for assistance with ultracentrifugation; and J.Q. Wu, T. Pollard, and D. Kellogg for sharing reagents and strains. This work was supported by grants from the NIH (R01 GM099774) and the American Cancer Society (RSG-15-140-01-CCG) to J.B.M, a shared instrumentation grant from the NIH (1S10OD018046), an NIH Training Grant (T32GM008704) to C.A.H.A, and the Dartmouth ASURE program for supporting U.M.

